# Modeling Research Project Grant Success Rates from NIH Appropriation History: Extension to 2020

**DOI:** 10.1101/2020.11.25.398339

**Authors:** Jeremy M. Berg

## Abstract

Communication of the likelihood that a grant proposal is funded is important for individual researchers and academic institutions. For research project grants (RPGs) funded by the US National Institutes of Health (NIH), this is measured by the success rate, the fraction of grants reviewed in a given fiscal year that are funded. In 2016, I posted a relatively simple computational model that allowed estimation of these success rates from historical data of NIH appropriations and associated data, based largely on the fact that commitments to fund NIH grants are typically made over four fiscal years rather than coming from a single fiscal year’s appropriations. Data for appropriations and success rates for fiscal years 2016-2020 are now available, allowing this model to be tested without no adjustable parameters. Over this period, the NIH appropriation increased by 29.7% to $39.313 billion, yet the RPG success rate increased from 0.183 to 0.201, a relative increase of only 9.8%. This difference is accurately reproduced by the model, indicating that the modest changes in success rate despite large appropriation increases reflect the consequences of the funding grants from multiple fiscal years and the response of the biomedical research community in submitting an increased number of proposals. The model can also be extended to estimate the success rates for individual institutes with reasonable accuracy.

## INTRODUCTION

A substantial portion of scientific research is government-supported. An important factor that affects the experiences of individual investigators and the efficiency of government funding programs is the likelihood that a given grant application is awarded funding. While this can seem to be a simple parameter, it depends on the systems properties of the scientific enterprise such as the fraction of appropriated funds available for new grants and the number of investigators and associated grant proposals competing for funding, both of which evolve over time.

Having served as Director of the National Institute of General Medical Sciences at NIH from 2003-2011, I experienced the consequences of this phenomenon directly, particularly since my service began at the end of the period when the NIH appropriation was approximately doubled from 1998-2003. In the last year of the “doubling” from fiscal year 2002 to fiscal year 2003, the NIH appropriation increased by 16.5% and the RPG success rate fell slightly from 0.312 to 0.299. In the next year, the appropriation increased by 3.8% and the RPG success rate fell dramatically to 0.246, a relative drop of 18%. This drop and those in subsequent years led to outcries from the biomedical research community, including criticisms of NIH priorities without an understanding that this large drop was essentially inevitable.

The most important contributors to these decreases in success rate relates to the financial structure of how NIH grants are funded. In almost all cases when a multi-year NIH grant is made, funds for the first year are provided from the appropriation for that fiscal year with commitments made to support additional years of the grant out of appropriations for subsequent years. With an average duration of four years, these commitments last for three additional years, at which point the funds are available for supporting competing grants. Because of this, 75% or more of each year’s appropriation is already committed with the remaining 25% or less, adjusted by any increase or decrease in the appropriation, being available for new or competing grants. This recycling process is shown in Figure 1.

**Figure 1.**
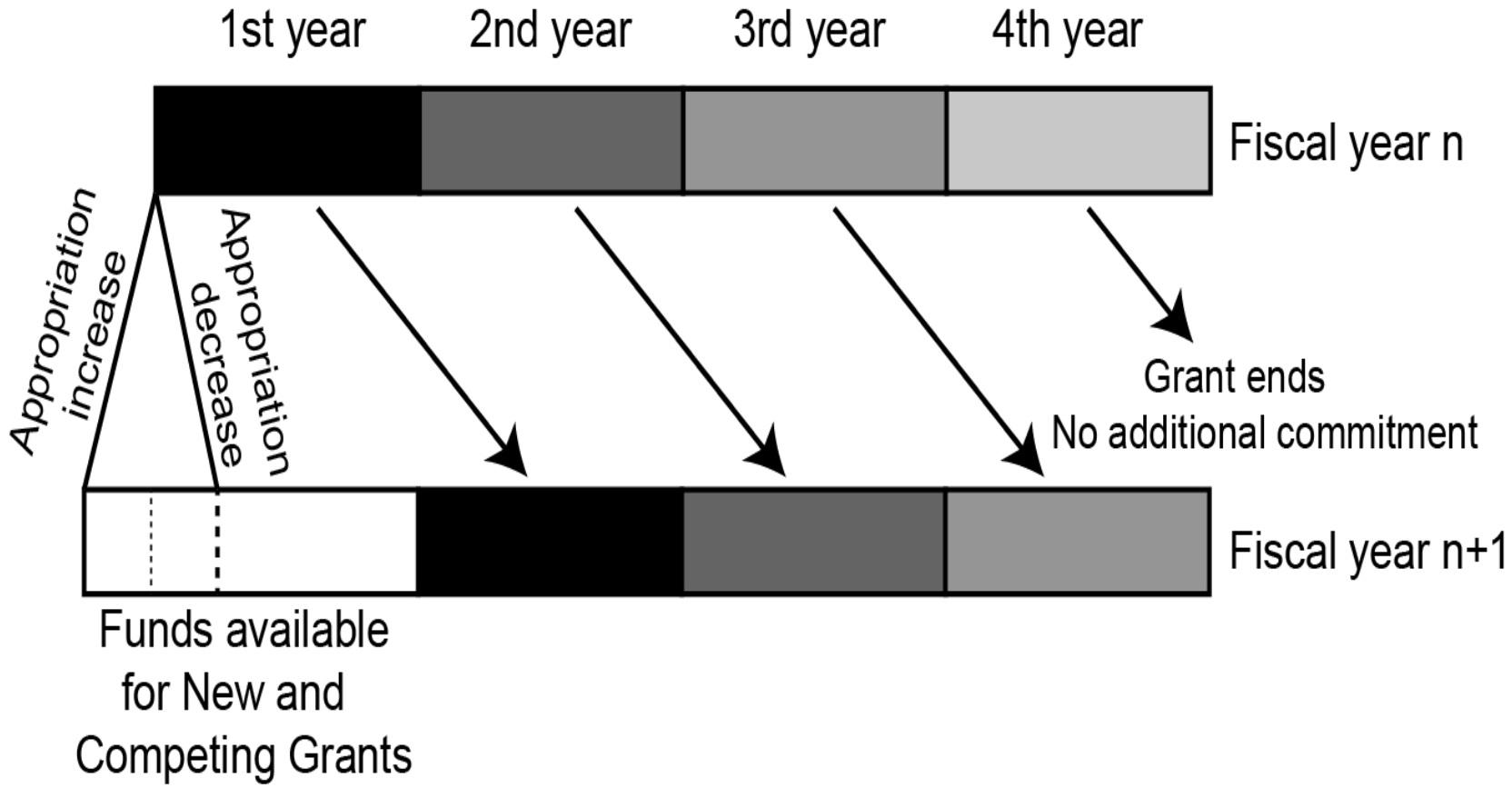
An illustration of how funding a four-year grant results in outyear commitments with funds only becoming available after the fourth fiscal year. The amount of these “recycled” funds is affected by any increase or decrease in the NIH appropriation.

This process amplifies the impact of any change in the appropriation by four-fold or more.

## RESULTS THROUGH 2015

I developed a computational model based on these processes and posted the result in 2016 including the underlying data and R code (https://blogs.sciencemag.org/sciencehound/2016/08/25/modeling-success-rates/, https://blogs.sciencemag.org/sciencehound/2016/09/29/modeling-the-annual-number-of-nih-research-grant-applications/). The estimation of the RPG success rate is based on two components: (1) Estimation of the number of new and competing RPGs based on the appropriation history and resource recycling; and (2) Estimation of the number of grant applications based on the observation that the number of applications approximately tracks the increase in NIH appropriations with a delay of approximately two years.

The results are shown in Figure 2A–C.

**Figure 2A.**
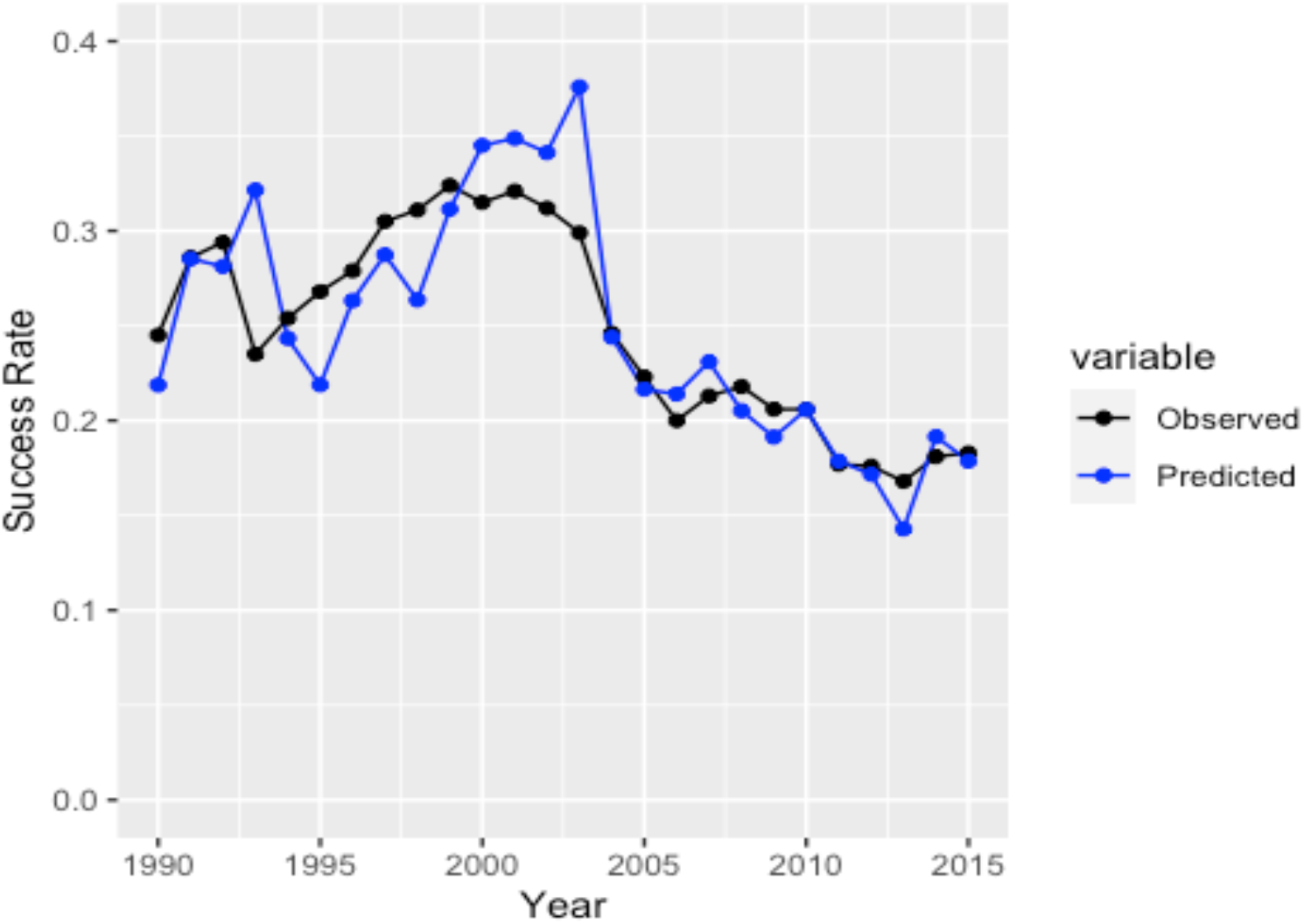
A comparison of observed and predicted success rates from 1990-2015 based on the previously reported model.

**Figure 2B.**
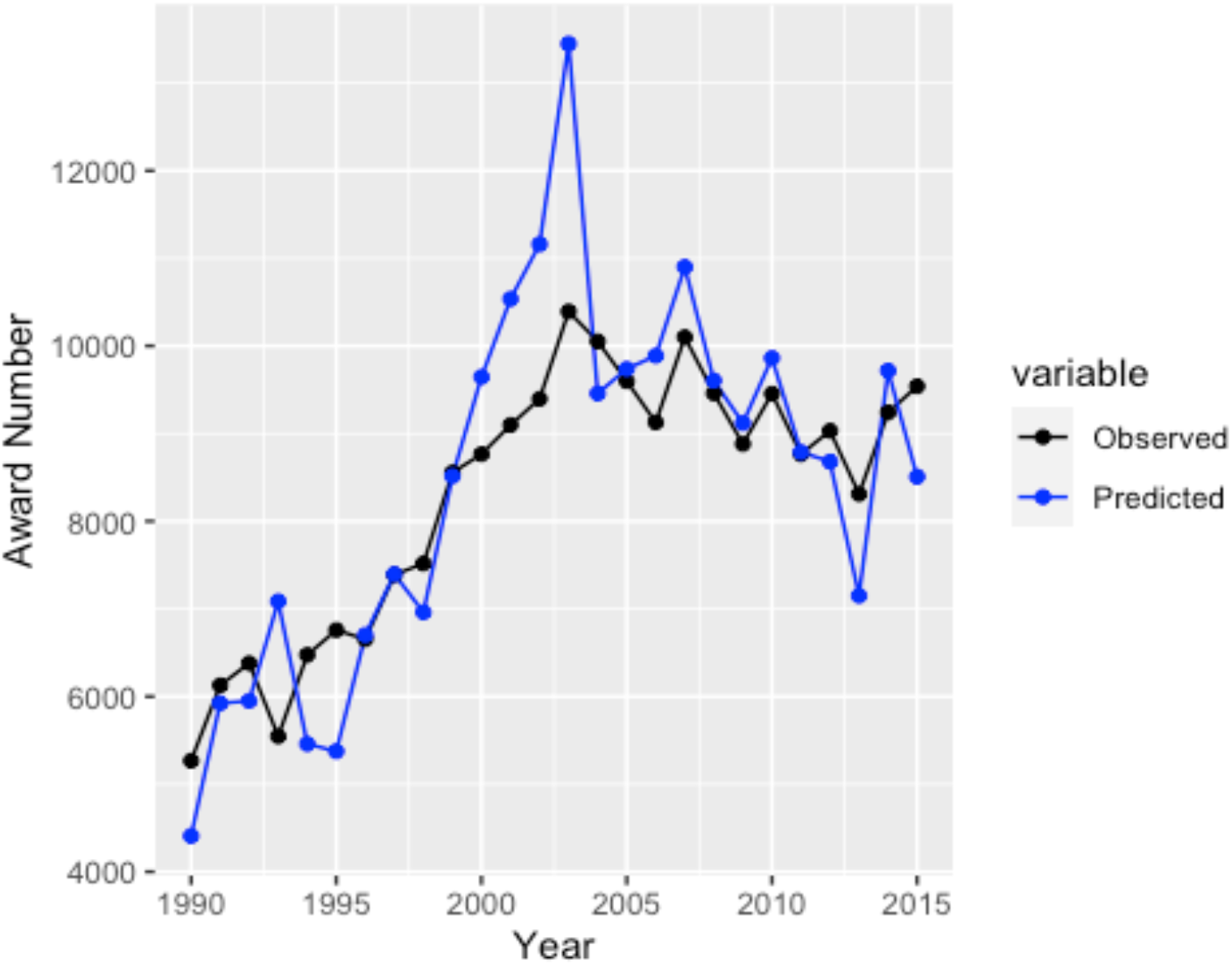
A comparison of observed and predicted number of competing RPG awards from 1990-2015 based on the previously reported model.

**Figure 2C.**
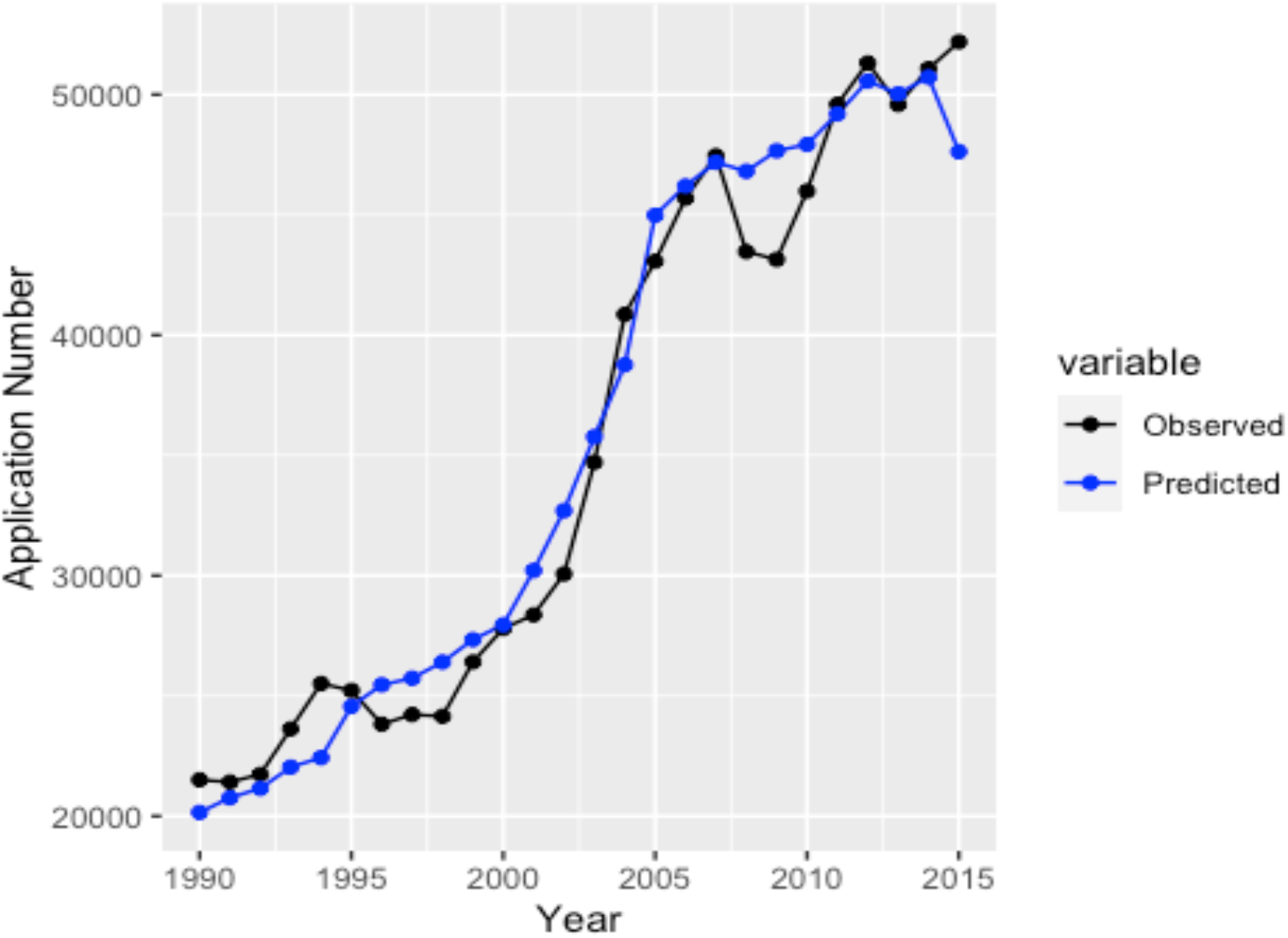
A comparison of observed and predicted number of competing RPG applications from 1990-2015 based on the previously reported model. The decreased number of applications in 2008 and 2009 is due to the fact that applications associated with the American Recovery and Reinvestment Act were not included. The number of applications is predicted based on the appropriation levels for the preceding two years using a regression model.

In general, the agreement is quite good. The largest differences are observed between 2001 and 2003 where the model predicts that more RPGs would have been funded than actually were. This reflects changes in NIH policies where non-RPG mechanisms were used to fund larger research efforts made possible by the substantial increases in the NIH appropriation.

## RESULTS FOR 2016-2020

Since appropriation data are the only necessary input for the RPG success rate model with no adjustable parameters, the availability of data for fiscal years 2016-2020 allows extension of the model to cover these years. Four years have passed since the posting of this model with appropriation data available for five years and the numbers of RPG awards available for four. The NIH appropriation history including these years is shown in Figure 3.

**Figure 3.**
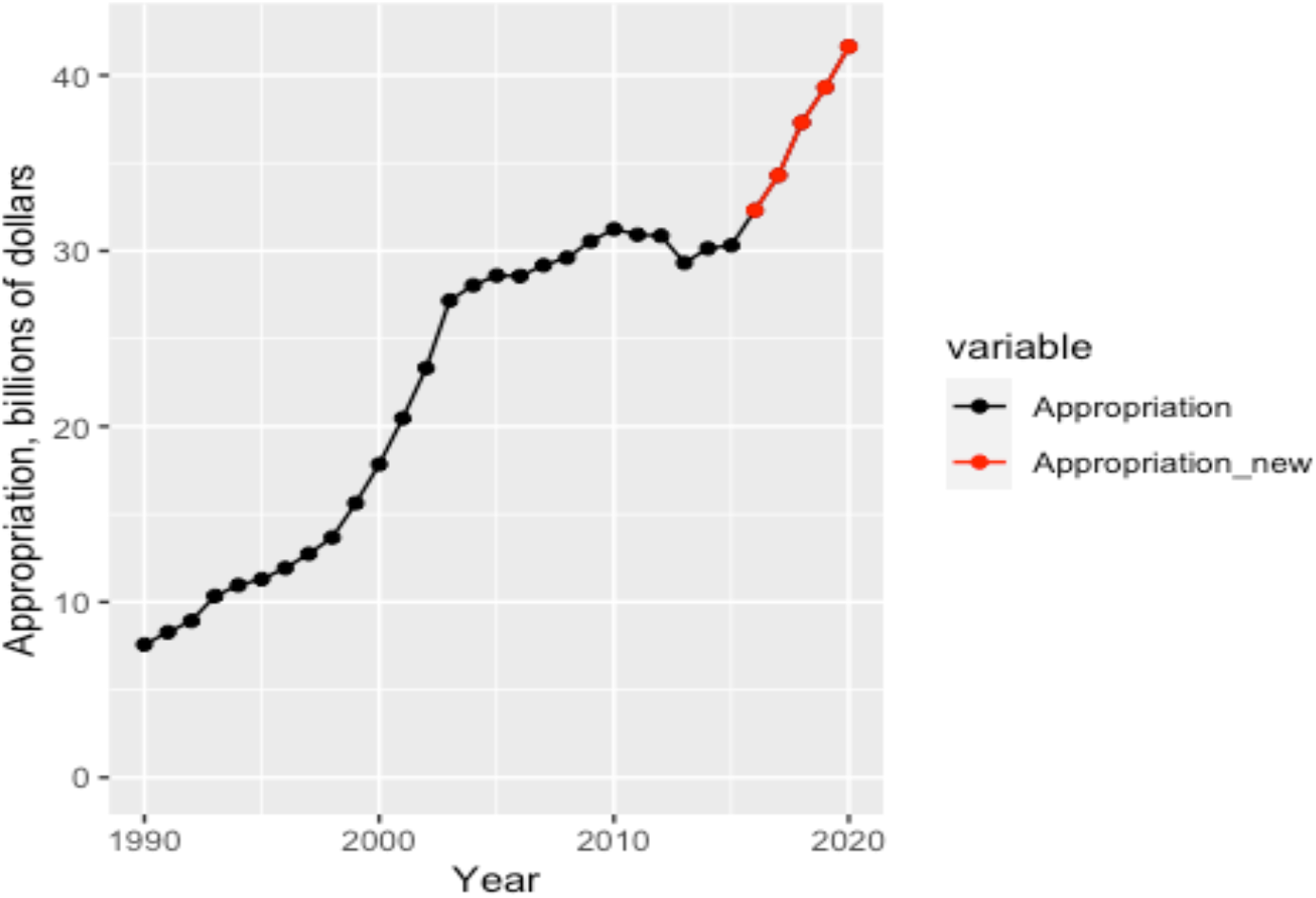
NIH Appropriations from 1990 to 2020. The five years (2016-2020) that occurred after the posting of the model, shown in red, reveal substantial increases in each year. The values shown are in nominal dollars, not corrected for inflation.

Success rate data are available for fiscal year 2016-2019 with those data for 2020 expected to be available in December, 2020. The results are shown in Figure 4.

**Figure 4.**
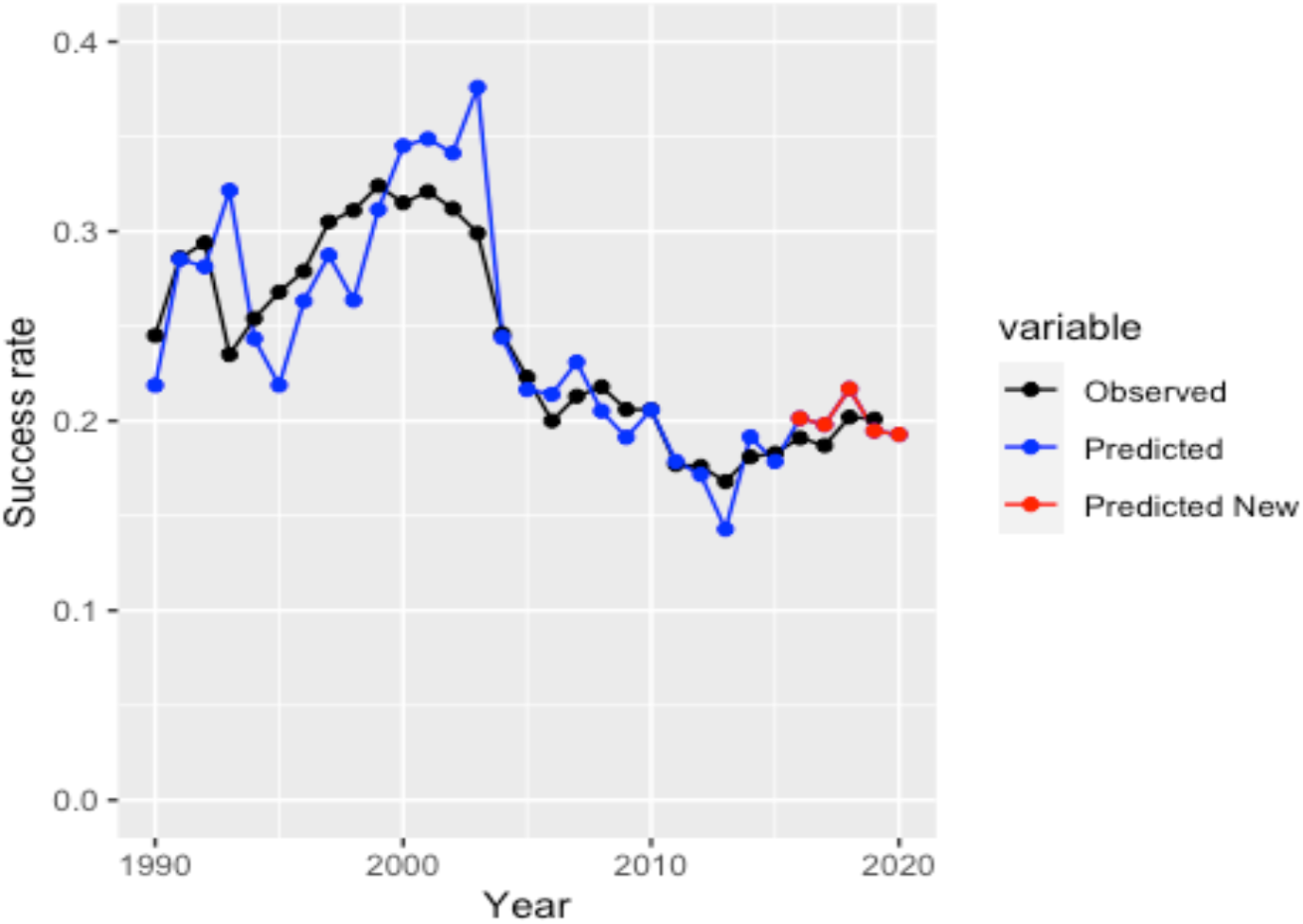
A comparison of the predicted and observed success rates including years 2016-2019 with the predicted value for 2020.

The agreement is reasonably good. The predicted success rates are slightly higher than the ones for 2016-2018 in a manner that is similar to, although less pronounced than, what was observed during the 2000-2004 period during the “doubling”. For 2016-2018, the predicted success rate exceeds the observed one by an average of 0.012, corresponding to a relative difference of 0.012/0.193, or 6.3%. For 2000-2004, the predicted success rate exceeds the observed one by an average of 0.032, corresponding to a relative difference of 0.032/0.299, or 10.9%.

The predicted and observed numbers of RPGs and numbers of applications are shown in Figures 5 and 6, respectively.

**Figure 5.**
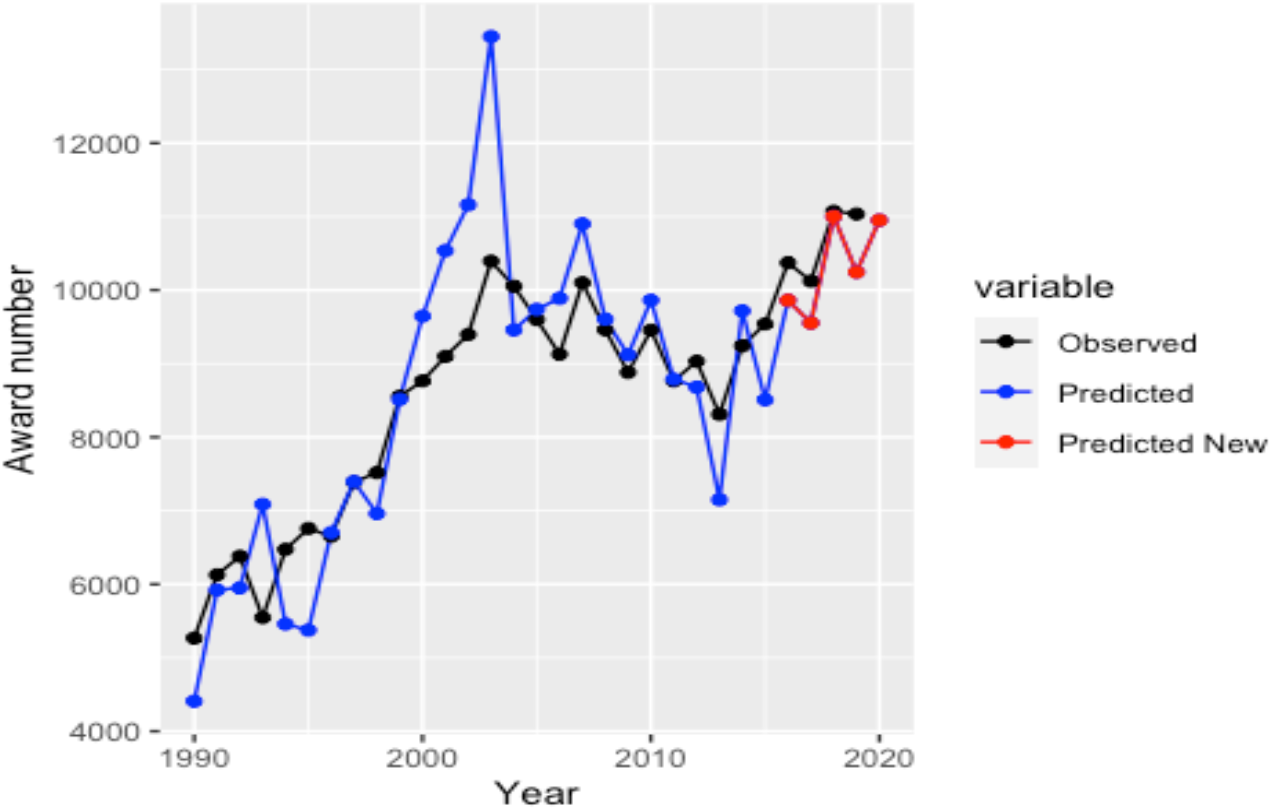
A comparison and observed numbers of RPG awards including years 2016-2020.

**Figure 6.**
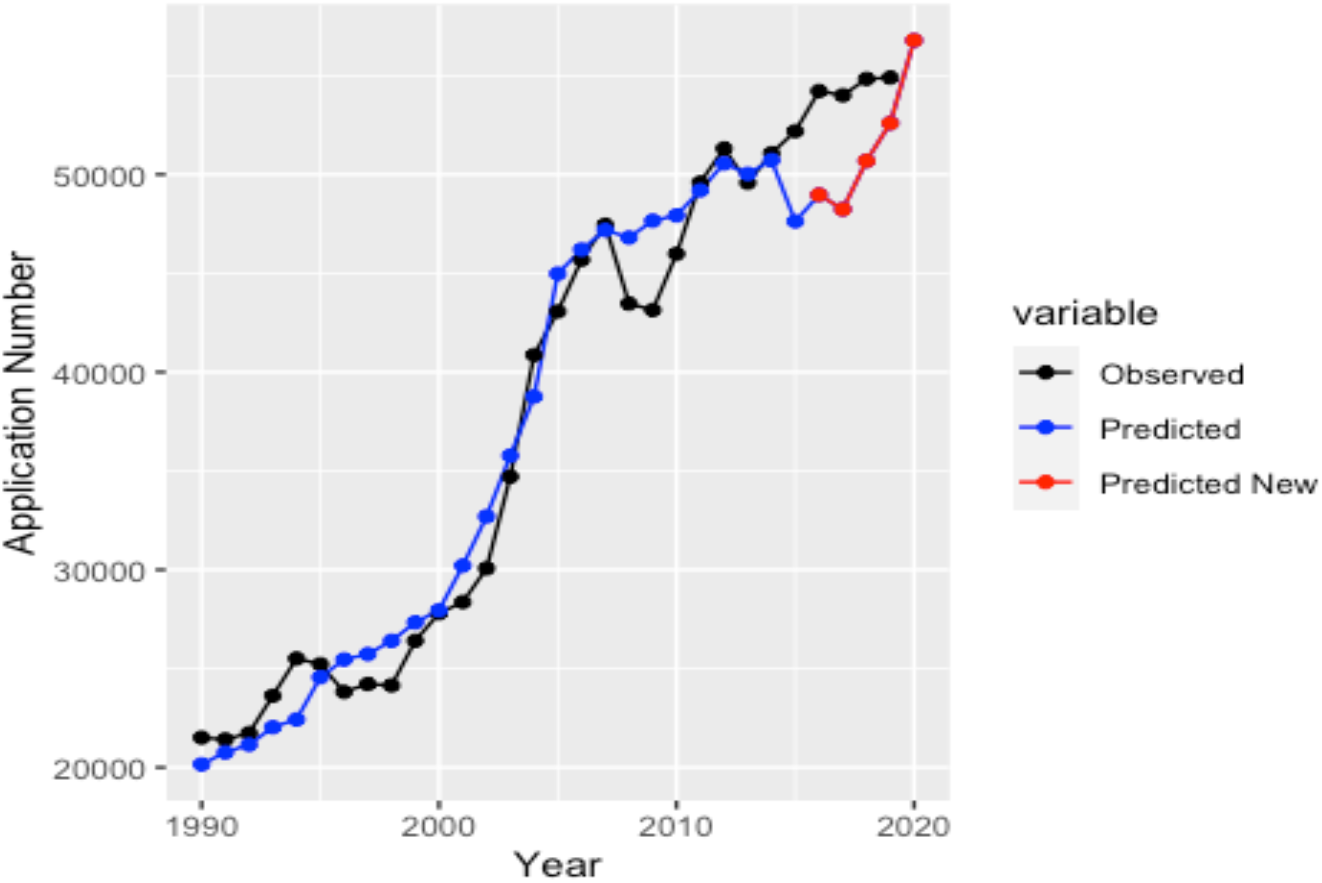
A comparison and observed numbers of RPG applications including years 2016-2020.

Figure 5 reveals that the model tracks the number of RPG awards fairly closely but the predicted number is lower than the observed number, unlike what happened during the “doubling”. This can be accounted for by some modest policy changes by NIH. From 2004 to 2015, the percentage of the overall NIH appropriation invested in RPGs averaged 51.5% with a standard deviation of 0.7%. From 2015 to 2019, this increased to 54.9%. Some of this increase went to increasing the average size of an RPG award from annual total costs of $477 thousand to $554 thousand but the remainder allowed for an increase in the number of awards. Note that larger increases in the percentage of the NIH appropriation devoted to RPGs and in the average size of an RPG also occurred during the doubling, but these changes were not sufficiently large to consume the annual increases of approximately 14% in the appropriation during the doubling, compared with approximately 7% from 2016 to 2020.

Figure 6 demonstrates that the number of RPG applications is also underestimated by the model. This difference is largest for 2015 where a substantial decrease in the number of applications is predicted. This is due to the unprecedented 5% decrease in the NIH appropriation from 2012 to 2013. This decrease in the number of applications is likely unreasonable since both appropriation increases (presenting additional opportunities) and appropriation decreases (implying increased competition for funding) likely lead to increases in proposal submissions. The predicted number of applications moves closer to be observed values in 2018 and 2019 and will likely exceed the observed value for 2020.

Since both the number of RPG awards and of applications are underestimated by the model, the success rate, the ratio of these quantities is more accurately predicted that either component. It is gratifying that the model with a least squares error of 0.011, corresponding to a relative error of 5.7% with no adjustable parameters. This compares with a least squares error of 0.03, corresponding to a relative error of 12.3% for the period from 1990 to 2015 over which the model was initially developed.

## EXTENSION TO R01 SUCCESS RATES FOR INDIVIDUAL NIH INSTITUTES AND CENTERS

While the overall NIH RPG success rate is of general interest, success rates for individual NIH institutes and centers for particular funding mechanisms such as R01 grant are likely to be more relevant to specific, individual investigators. Institute- and center-specific R01 success rates are readily available for year 2001 and later.

The NIH-wide R01 success rate for 2001-2019 is highly correlated with the NIH-wide RPG success rate with a Pearson correlation coefficient of 0.986. The R01 success rates for individual institutes and centers are generally well correlated with the NIH R01 success rates with 4 institutes with correlation coefficients > 0.9 (NIDCD, NHLBI, NIAID, NIDA), 8 additional institutes with correlation coefficients > 0.8 (NINDS, NCI, NIGMS, NIDDK, NIDCR, NICHD, NIEHS, NIA), and 5 additional institutes with correlation coefficients > 0.7 (NIMH, NIAMS, NINR, NEI, NIAAA). These institutes are those that have relatively large appropriations with substantial investments in R01s while those with lower correlation coefficients have smaller appropriations or prioritize other funding mechanisms. Taken together, the 17 institutes with correlation coefficients > 0.7 account for 87.1 % of the overall NIH appropriation for fiscal year 2020.

Based on these correlations, the R01 success rates for these individual institutes can be estimated from the predicted RPG success rate by fitting each R01 success rate to the predicted RPG success rate using linear models. This approach can be reasonably successful as shown with the data for NIGMS shown in Figure 7.

**Figure 7.**
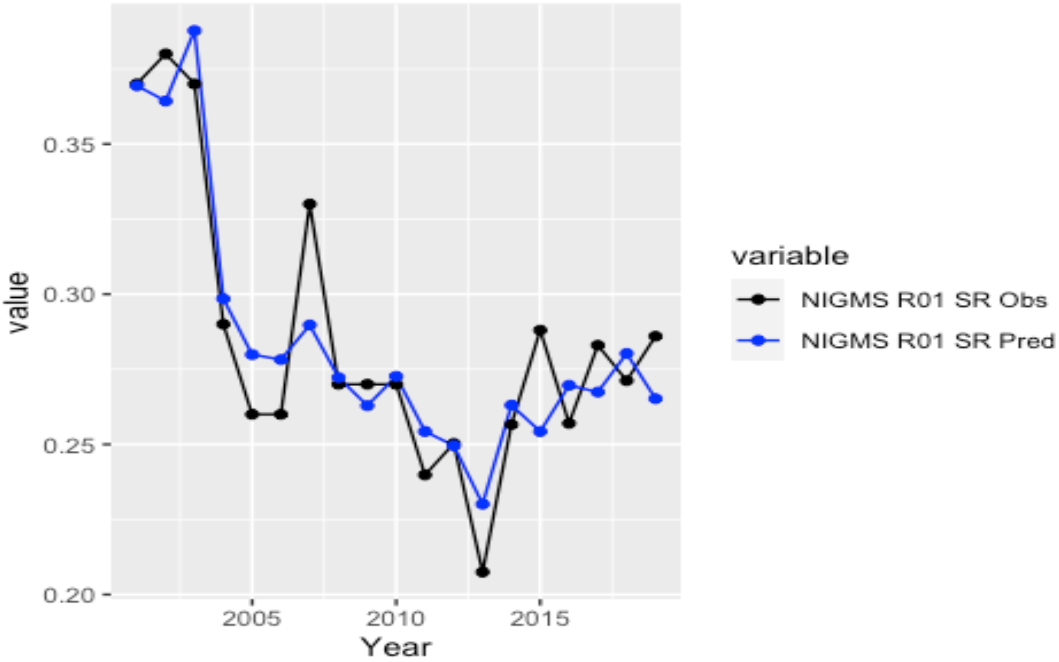
A comparison of the observed and predicted R01 success rates for NIGMS. The predicted success rates are derived from the predicted NIH-wide RPG success rate using a linear regression model.

The agreement is reasonably good across the entire period from 2001 through 2019 with a root mean squared deviation of 0.018.

Some of the limitations of the use of a simple linear regression model for different institutes are revealed with the data from the National Cancer Institute (NCI), shown in Figure 8.

**Figure 8.**
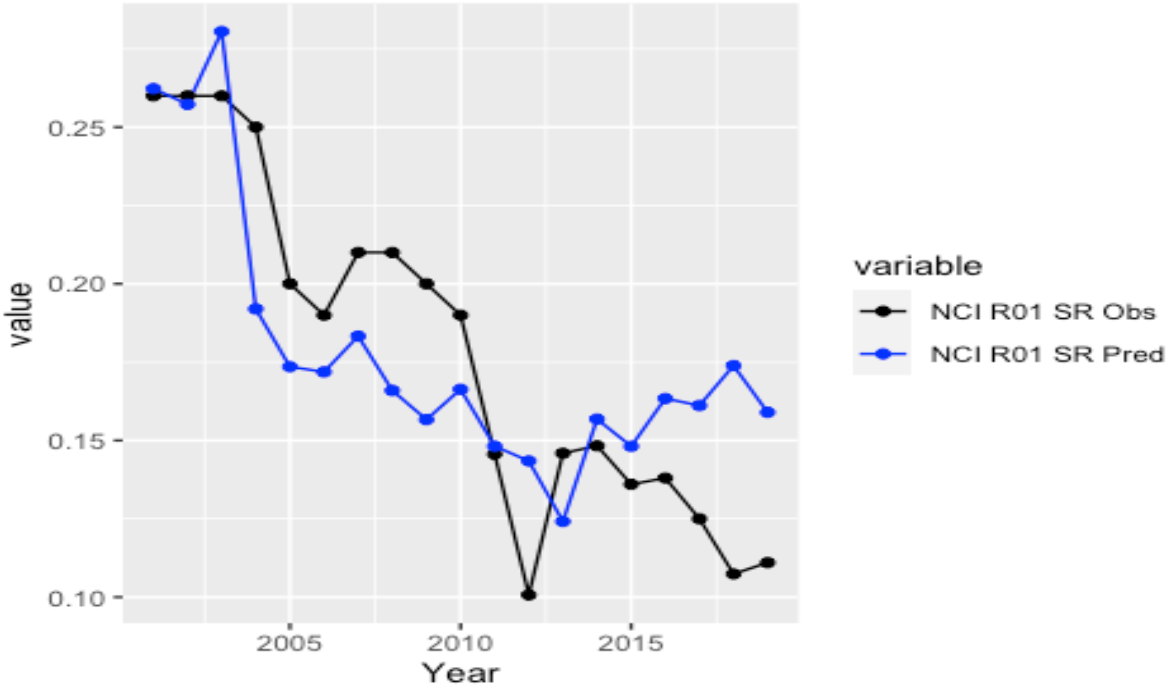
A comparison of the observed R01 success rates for NCI using a single, linear regression model from the predicted NIH-wide RPG success rate.

Figure 8 reveals that the model captures some aspects of the R01 success rate trend, but the observed and predicted R01 success rates diverge from 2016 to 2019. The agreement across the entire period from 2001 through 2019 with a root mean squared deviation of 0.033. The point of this divergence coincides with the launching of Cancer Moonshot (https://www.cancer.gov/news-events/press-releases/2016/ncab-nci-accept-brp-report), a major effort to accelerate the pace of cancer research. While new resources were committed to Cancer Moonshot project, the focus on this effort also led to an increase in applications. From 2015 to 2019, the number of R01 applications assigned to NCI increased by 30%, compared to an increase of 11% across NIH. The number of NCI R01 awards increased by 13% over this period. The substantial increase in applications over awards drove the R01 success rate down.

In order to account for this change in behavior, the success rate prediction regression can be done segmentally with one segment from 2001 to 2014 and a second from 2015 to 2019. The result is shown in Figure 9.

**Figure 9.**
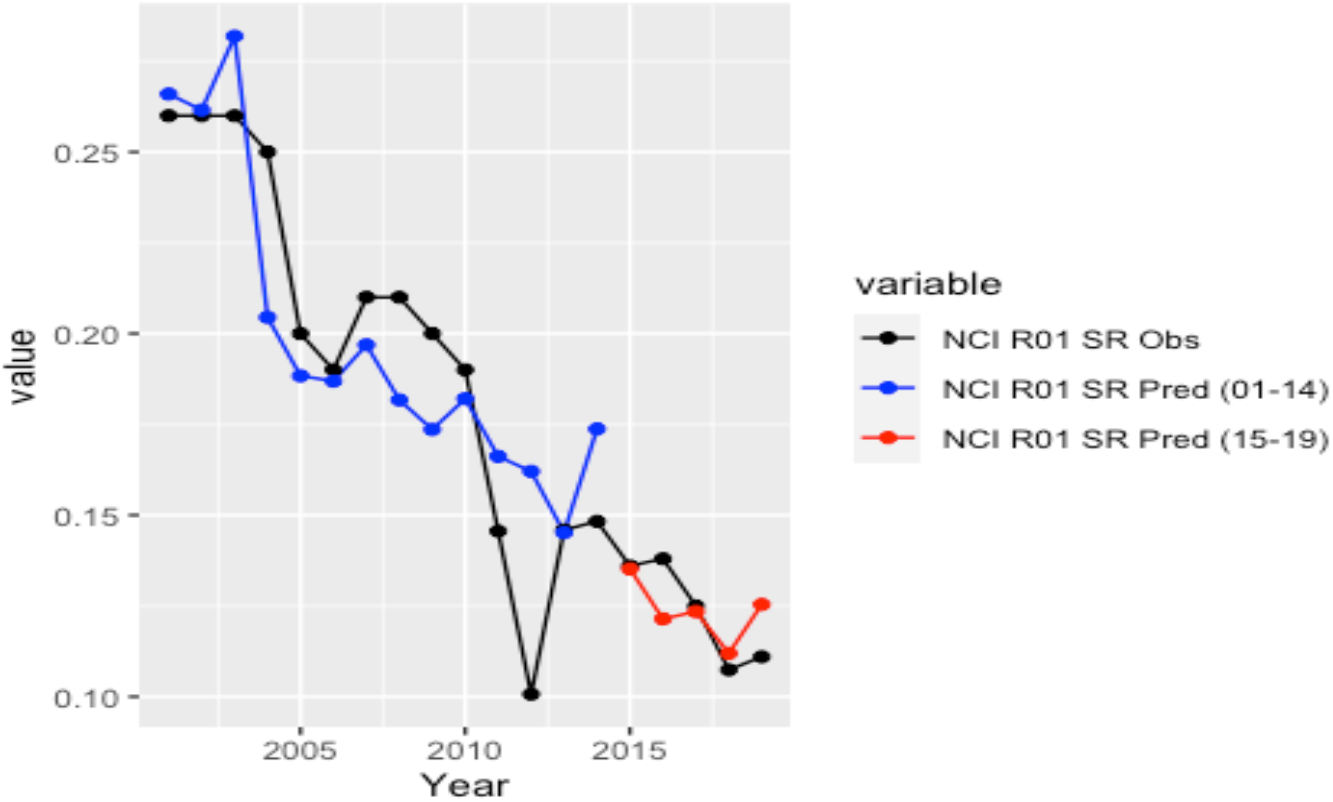
Modeling the R01 success rate for NCI using linear regression models over two periods: 2001 to 2014 and 2015 to 2019.

This approach enables the alignment of the predicted and observed R01 success rates across the entire period and reduces the root mean squared deviation to 0.023 from 0.033. This illustrates some of the challenges of estimating success rates for individual institutes and centers and one approach to improve the predictions through the addition of other information. However, for simplicity, only single linear regression models will be used for all institutes in all further analysis.

A final parameter of interest is the relative R01 success rates for each institute and center compared to the overall NIH-wide success rate. These ratios are shown in Figure 10.

**Figure 10.**
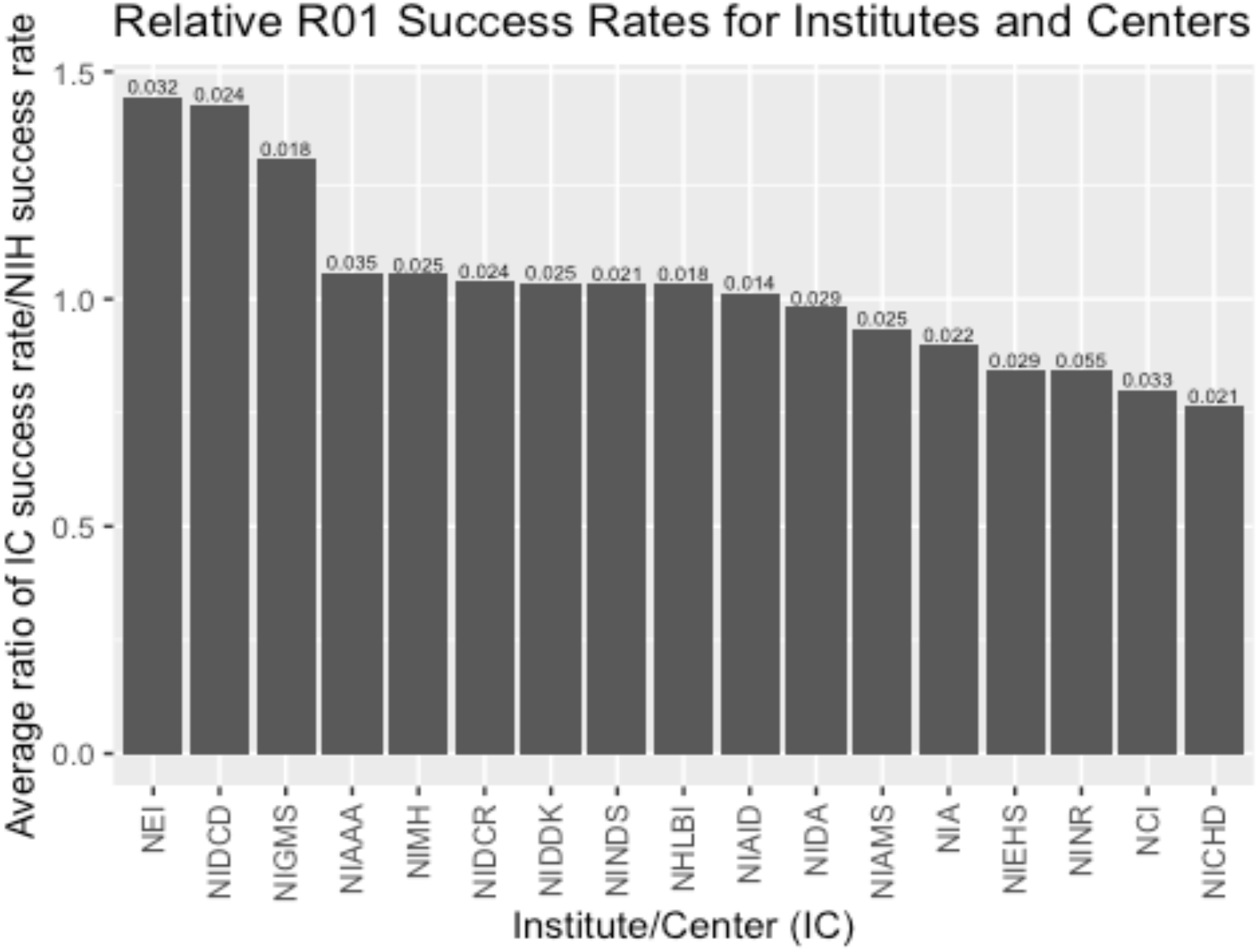
The ratio of R01 success rates for all NIH institutes with correlation coefficients with the NIH-wide R01 success rate > 0.7 averaged over the period from 2001 to 2019. The root mean squared error between the observed and predicted R01 success rates is shown over the bar for each institute.

This reveals that the R01 success rates are substantially higher than the NIH-wide average for three institutes: The National Eye Institute (NEI), the National Institute of Deafness and other Communication Disorders (NIDCD), and NIGMS. Six institutes show average R01 success rates lower than the NIH-wide value with the three lowest being the National Institute of Nursing Research (NINR), NCI, and the Eunice Kennedy Shriver National Institute on Child Health and Human Development. The deviations from the NIH-wide averge reflect institutes that have more R01 applications assigned to them then can be supported by their appropriations or that favor the use of other mechanisms over R01 grants.

Taken together, these results demonstrate that a simple model that captures the essence of the effects of multi-year grant funding but does not require more detailed information, can be used to estimate annual RPG success rates across NIH with surprisingly good accuracy. Moreover, these NIH-wide RPG success rates can be extended to estimate R01 success rates for many institutes. These models can be used to provide guidance for these success rates once the overall NIH appropriation is available.

## METHODS

Data for NIH appropriations were from https://officeofbudget.od.nih.gov/approp_hist.html and for numbers of RPG applications and success rates from https://report.nih.gov. Data for the Biomedical Research and Development Price Index (BRDPI) were from https://officeofbudget.od.nih.gov/pdfs/FY21/gbi/BRDPI_FY2019_2025_Final.pdf.

The model for the number of RPG awards is based on the following assumptions:

1. The average RPG has a duration of 4 years with 1/4 with a duration of 3 years, 1/2 with a duration of 4 years, and 1/4 having a duration of 5 years.
2. 50% of the NIH appropriation each year is devoted to supporting RPG awards.
3. The average RPG award size is increased for inflation using the BRDPI value from the preceeding year.

Each year, the amount required for all that have not passed their final year are subtracts from the RPG allotment from the NIH appropriation. The remaining funds are divided by the updated average grant size to yield the number of RPG awards that can be made. The funds are divided between the pools of 3-year, 4-year, and 5-year awards.

The model was intialized in 1980 with a pool of 17,000 RPGs awards with an average size of $100,851, distribute equally among the years of their duration. The model was allowed to run until 1990 to allow the distribution of awards to approximate the distribution of awards are various stages. The model was then extended from 1990 to 2020.

The model for the number of RPG applications is based on the observation that the shape of the number of applications over time approximately follows that for the increase in appropriation with a delay of approximately two years. The number of RPG applications was estimated by fitting the fractional change in applications (normalized to 1990) to the expression

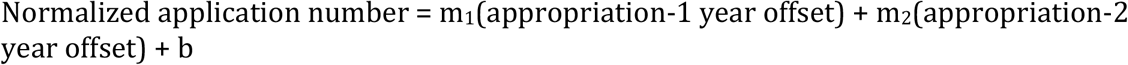

where appropriation-1 year offset is the normalized appropriation from the preceding year, appropriation-2 year offset is the normalized appropriation from the year two years prior, and b is a constant.

Linear regression using data from 1990 to 2015 yielded m_1_ = −0.18, m_2_ = 0.61, and b = 0.57. The agreement between the predicted and observed application numbers was good with a Pearson correlation coefficient of 0.983.

The predicted success rate was calculated by dividing the predicted number of RPG awards by the predicted number of applications.

R01 success rates for individual NIH institutes and centers were estimated using linear regression models including a constant term.

## Notes

https://officeofbudget.od.nih.gov/approp_hist.html

https://officeofbudget.od.nih.gov/pdfs/FY21/gbi/BRDPI_FY2019_2025_Final.pdf

https://blogs.sciencemag.org/sciencehound/2016/08/25/modeling-success-rates/

https://blogs.sciencemag.org/sciencehound/2016/09/29/modeling-the-annual-number-of-nih-research-grant-applications/

### Competing Interest Statement

The authors have declared no competing interest.

